# Functional interaction between Fanconi anemia and mTOR pathways during stalled replication fork recovery

**DOI:** 10.1101/2020.01.16.899211

**Authors:** Matthew Nolan, Kenneth Knudson, Marina K. Holz, Indrajit Chaudhury

**Affiliations:** University of Minnesota, Morris, MN; New York Medical College, Valhalla, NY; presently at St. Mary’s College of Maryland, St. Mary’s City, MD

## Abstract

We have previously demonstrated that Fanconi Anemia (FA) proteins work in concert with other FA and non-FA proteins to mediate stalled replication fork restart. Previous studies suggest a connection between FA protein FANCD2 and a non-FA protein mechanistic target of rapamycin (mTOR). A recent study showed that mTOR is involved in actin-dependent DNA replication fork restart, suggesting possible roles in FA DNA repair pathway. In this study, we demonstrate that during replication stress mTOR interacts and cooperates with FANCD2 to provide cellular stability, mediates stalled replication fork restart and prevents nucleolytic degradation of the nascent DNA strands. Taken together, this study unravels a novel functional cross-talk between two important mechanisms: mTOR and FA DNA repair pathways that ensure genomic stability.

## INTRODUCTION

Fanconi Anemia (FA) is an inherited genomic instability disorder characterized by predisposition to a variety of cancers, most notably acute myeloid leukemia and solid tumors. At least 22 FA proteins are known to function in a network to perform their genomic caretaker role [1]. FA pathway proteins have been extensively studied for their roles in the repair of a lethal type of DNA damage called DNA inter-strand crosslinks (DNA ICLs) [2–6]. However, in the past several years, additional functions of FA pathway proteins were discovered [7]. One such function is to mediate stalled DNA replication fork restart [8–10]. Upon insult from replication inhibitors, ongoing replication fork stalls. FA protein FANCD2, along with other FA [10] and non-FA proteins [8, 9, 11, 12] work in a common pathway that ensures that, once the replication inhibitor is removed, the stalled fork can resume DNA replication, which is essential for the completion of genome duplication.

In addition to replication fork restart, FANCD2 mediates another function at the stalled fork. During fork stalling, nascent DNA strands at the stalled fork are vulnerable to nucleolytic degradation by cellular nuclease such as Mre11. FANCD2, along with breast cancer associated proteins BRCA1 and BRCA2 (also known as FA proteins FANCS and FANCD1, respectively), prevent Mre11 from degrading the nascent DNA strands, and thus contribute to the fork stability [8, 9, 13].

A non-FA protein, mechanistic target of rapamycin (mTOR) regulates central cellular functions, such as proliferation, growth, viability, transcription and translation, and autophagy [14–16]. mTOR has an important role in cancer, and deregulation of mTOR pathway components has been implicated in multiple cancer types [14, 17–21]. Thus, inhibition of mTOR pathway has become a viable therapeutic strategy, and mTOR inhibitors are FDA-approved for several oncology indications [15, 16, 18, 20, 22–25]. mTOR has been shown to act in two complexes with divergent functions, mTOR Complex 1 and mTOR Complex 2 (mTORC1 and mTORC2) [26]. mTORC1 is acutely sensitive to an allosteric inhibitor rapamycin, and is the focus of this work.

In a recent study, mTOR was shown to be essential for stalled replication fork restart. mTOR works in a pathway that involves actin-dependent replication fork recovery, and inhibition of mTOR inhibited stalled replication fork restart [27]. This finding warrants further investigation to determine the roles of mTOR and the FA pathway during replication fork stalling. Interestingly, mTOR inhibition in human hematopoietic stem/progenitor (HSPC) cells was shown to cause phosphorylation and nuclear import of NF-κB, and subsequent binding of NF-κB to the promoter of FANCD2 which inhibits FANCD2 expression, connecting mTOR and the FA DNA damage response pathway [28]. In this study, we show evidence for the concerted actions of both mTOR and the FA pathway protein FANCD2. We demonstrate that mTOR and FANCD2 are essential and work in the same pathway to mediate aphidicolin- (APH; inhibitor of DNA polymerase alpha that causes fork stalling) induced stalled replication fork restart and provide stability of the nascent DNA strands from nucleolytic degradation at the stalled forks.

## RESULTS

### mTOR, in concert with FANCD2 maintains cellular survival in response to replication fork stalling agents

In order to determine if there is a functional cross-talk between mTOR and FA pathways in response to DNA damage or replication inhibition, we performed a cellular viability assay using FANCD2-deficient and proficient cells that were treated with mTOR inhibitor rapamycin to create mTOR-inhibited and FANCD2/mTOR double-deficient cells. These cells were next treated with the indicated doses of aphidicolin (APH, a replication inhibitor [29], Fig. 1A), hydroxyurea (HU, a replication inhibitor [30], Fig. 1B) and mitomycin C (MMC; DNA interstrand crosslinking agent [31], Fig. 1C). As seen in Figure 1, APH treatment significantly reduced cellular viability of the FANCD2-deficient and mTOR-inhibited cells, while no additive effect was observed in the double-deficient cells, suggesting that FANCD2 and mTOR are epistatic for providing cellular survival in response to APH (Fig. 1A). A similar but somewhat milder effect was visible in the mTOR-inhibited cells in response to HU treatment, suggesting that mTOR cooperates with FANCD2 in ensuring cellular viability (Fig. 1B). In response to MMC, however, mTOR-inhibited cells did not exhibit reduced cellular viability compared to that of the control, suggesting that mTOR is not important for providing cellular resistance to MMC induced DNA-ICLs (Fig. 1C).

**Fig. 1.**
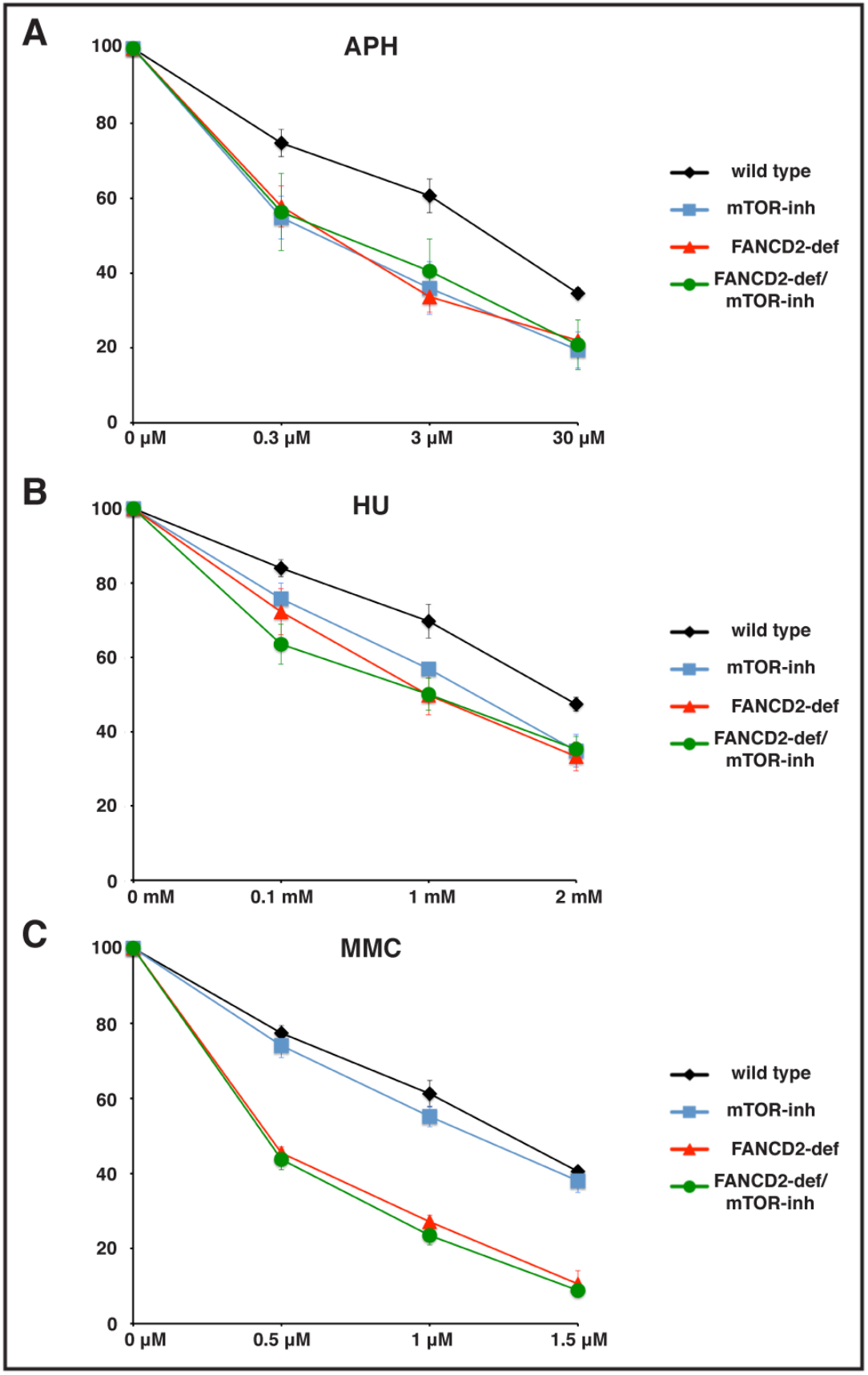
mTOR is essential, and acts in the same pathway with FANCD2 to maintain cellular viability during replication stress. FANCD2 deficient patient fibroblast cells (PD20) and PD20 cells retrovirally complemented with wild type FANCD2 (PD20+D2, wild type) were either untreated or treated with the mTOR inhibitor rapamycin (20 nM) to create mTOR-inhibited and FANCD2/mTOR double deficient cells. 500 cells from each condition were seeded on 60 mm dishes and allowed to grow for 8-10 hours. Cells were next treated with the indicated doses of APH, HU or MMC for 6 hours and allowed to grow in fresh media for 7-10 days. Colony forming units were counted after staining with crystal violet and results plotted as percentage normalized to untreated. Data plotted are means from three independent experiments.

### FANCD2 interacts with mTOR only in response to DNA damage or replication stress

Previous studies strongly suggest an interesting connection between FA protein FANCD2 and mTOR. mTOR was shown to regulate DNA damage through NF-κB-mediated FA pathway [28]. Using nuclear extracts of wild type (PD20+D2) fibroblast cells, we have performed immunoprecipitation (IP) assay with anti-FANCD2 antibody. As shown in Fig. 2A (i), FANCD2 specifically interacted with mTOR only after treatment with mitomycin C (MMC, a chemotherapeutic drug that induces DNA ICL), hydroxyurea (HU, replication inhibitor) or aphidicolin (APH, replication inhibitor) (Fig. 2A, lanes 10-12), but not in the untreated condition (Fig. 2A(i), lane 9). Quantitative analysis of FANCD2 and mTOR Western blot band intensities from three independent IP experiments (as shown in Fig. 2A(i) and in the inset) clearly suggests that mTOR interacts with FANCD2 only during DNA damage or replication stress induced by MMC, HU or APH but not in the untreated samples. Reciprocal IP with anti-mTOR antibody was performed using nuclear extracts from wild type (PD20+D2) and FANCD2-deficient cells (PD20) that were either untreated or treated with MMC or APH (Fig. 2B). FANCD2 preferentially interacted with mTOR during APH treatment (Fig. 2B, lane 9). FANCD2-mTOR interaction in response to MMC was significantly weaker (Fig. 2B, lane 8), and IP from untreated nuclear extract did not show any interaction (Fig. 2B, lane 7). These findings strongly suggest a possible functional connection between the FA and mTOR pathways during DNA damage and replication stress.

**Fig. 2.**
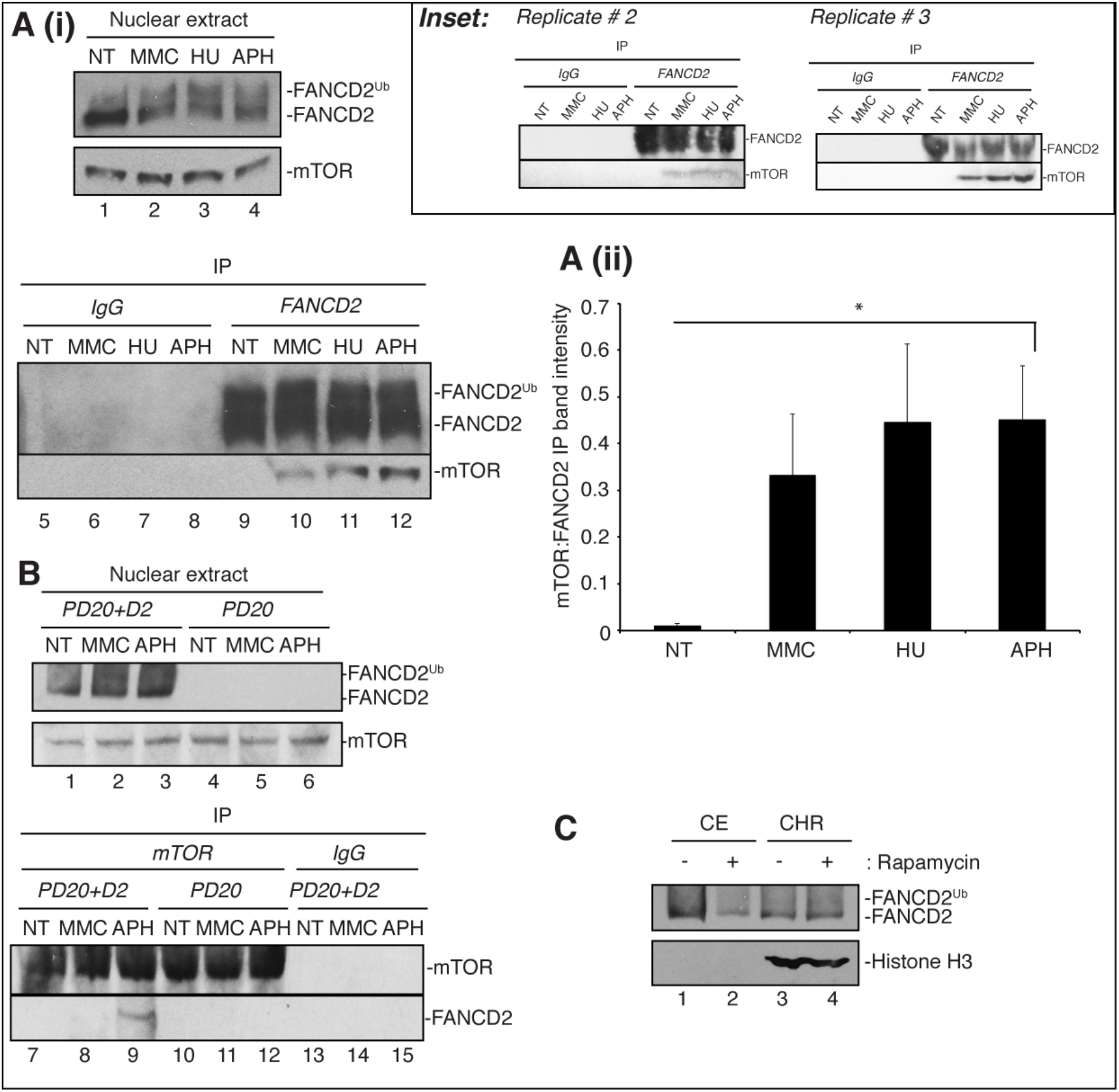
DNA damage induces FANCD2/mTOR interaction. (A (i)) A FANCD2-deficient patient fibroblast cell line (PD20) retrovirally complemented with wild type FANCD2 (PD20+D2) was used for this experiment. Cells were either untreated (NT) or treated with 1 μM MMC for 24h, 1 mM HU for 24h or 30 μM APH for 6 hours. Nuclear extracts were isolated (lanes 1-4) and subjected to immunoprecipitation (IP) using IgG (lanes 5-8) or anti-FANCD2 antibody (lanes 9-12). IP was performed in presence of ethidium bromide (10μg/ml), a DNA intercalating agent, to rule out the possibility of DNA-bridged interactions between these proteins. Nuclear extracts and IP samples were analyzed by Western blotting for the presence of FANCD2 and mTOR. Inset: Repetition of FANCD2-IP experiments. (A(ii)) Western blot band intensities from three independent IP experiments as shown in (A(i)) were measured by ImageJ software. Ratio of mTOR and FANCD2 IP bands (mTOR : FANCD2) from NT, MMC, HU and APH treated IP samples were plotted. Error bar represents standard error of mean. * denotes p value < 0.05. (B) Reciprocal IP was performed using nuclear extracts from wild type (PD20+D2; lanes 1-3) and FANCD2-deficient cells (PD20; lanes 4-6). Cells were either untreated (NT) or treated with 1 μM MMC for 24h or 30 μM APH for 6 hours. IP was performed using anti-mTOR antibody (lanes 7-12) or IgG (lanes 13-15). IP was carried out in presence of ethidium bromide (10μg/ml) to rule out the possibility of DNA-bridged interactions between these proteins. Nuclear extracts and IP samples were analyzed by Western blotting for the presence of FANCD2 and mTOR. (C) Cytoplasmic (lanes 1-2) and chromatin extracts (lanes 3-4) from wild type (PD20+D2) cells either untreated (lanes 1, 3) or treated with 20 nM mTOR inhibitor rapamycin (lanes 2, 4) reveal that although absence of mTOR activity reduced FANCD2 expression, FANCD2 chromatin recruitment was not affected.

### mTOR inhibition does not have any affect on FANCD2 chromatin recruitment, although it slightly down regulates FANCD2 expression

In order to determine if mTOR inhibition has any affect on FANCD2 expression and FANCD2 chromatin recruitment, we performed sub-cellular fractionation assay from untreated or rapamycin-treated wild type (PD20+D2) cells (Fig. 2C). While FANCD2 was enriched less in the rapamycin mediated mTOR inhibited cytoplasmic extracts compared to the untreated condition (Fig. 2C, lane 2 vs 1); no significant difference was observed in the chromatin recruitment of FANCD2 (Fig. 2C, lane 4 vs 3). This result suggests that although mTOR inhibition might have some effect in downregulating FANCD2 expression, FANCD2 chromatin recruitment is unaffected in response to mTOR inhibition.

### FA and mTOR pathways functionally interact to ensure efficient replication fork recovery

Our previous studies have shown that FA pathway protein FANCD2 is essential to mediate stalled replication fork restart [8, 9, 11, 12]. A recent study showed the importance of mTOR in mediating actin-dependent stalled fork restart [27]. As we have observed FANCD2 to directly interact with mTOR in response to replication stress (Fig. 2A and 2B) and FANCD2 and mTOR working together to provide cellular resistance to replication fork stalling agents (Fig. 1), we wanted to determine if mTOR is essential for FA-dependent stalled fork restart. To this end, we performed dual-labeling DNA fiber assay. As expected, untreated cells did not have any defect in replication fork restart, however, in response to APH, mTOR inhibition severely affected stalled fork restart compared to that of the wild type (31% in mTOR-inhibited cells vs 81% in control). FANCD2-deficient cells had a similar fork restart deficiency (37%). FANCD2/mTOR-double deficiency did not exacerbate the phenotype (29% fork restart), again supporting the notion that FANCD2 and mTOR are epistatic for stalled replication fork restart (Fig. 3C).

**Fig. 3.**
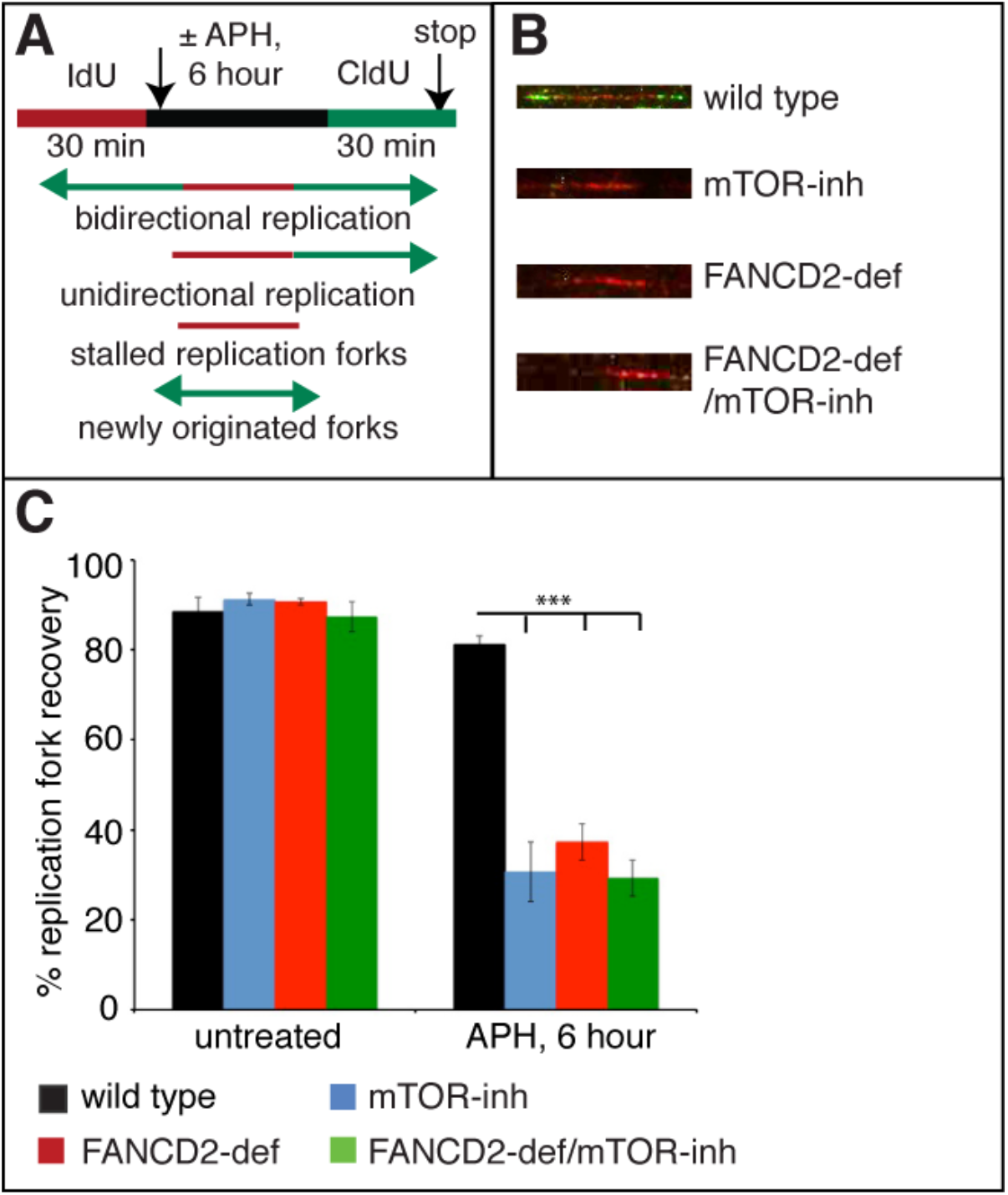
mTOR is essential for and works together with FANCD2 to mediate stalled replication fork restart. *(A) Schematics of DNA fiber assay.* FA patient fibroblast cells PD20 (FANCD2-deficient) and PD20 cells complemented with FANCD2 (PD20+D2, wild type) were treated with mTOR inhibitor rapamycin (20 nM) to create FANCD2/mTOR-double-deficient and mTOR-deficient cells respectively. For performing DNA fiber assay in these cells, 50 μM IdU were added to media and incubated for 30 minutes. Next, cells were either untreated or treated with 30 μM APH for 6 hours followed by incubation for 30 min in fresh media containing 50 μM CldU. DNA fibers were extracted, stretched on silane-coated slides, fixed and incubated with primary and fluorescent labeled secondary antibodies (Red color for IdU and green for CldU). Images were captured using a Deltavision microscope (University of Minnesota, Twin-Cities). Replication fork restart was measured by counting the proportion of restarted forks (Red-Green tract) compared to total forks (Red-Green + Red only tracts). *(B) Representative images of DNA fibers.* Representative images showing DNA fibers isolated from wild type, mTOR-inhibited, FANCD2-deficient, and FANCD2/mTOR-double deficient cells that were allowed to restart after APH stress. *(C) mTOR-inhibited cells have severe defect in the stalled replication fork restart similar to that of cells with FANCD2 deficiency.* DNA fiber assay was performed using wild type, mTOR-inhibited, FANCD2-deficient and FANCD2/mTOR-double deficient cells as shown in the schematics (A). Results plotted were from three independent experiments. Error bar represents standard error of mean. *** denotes p value ≤ 0.001.

### FA and mTOR pathways work together to ensure stability of the nascent DNA strands after replication fork stalling

Previous studies have established the role of FANCD2 in providing protection for replication forks from nucleolytic degradation by cellular nucleases such as Mre11, upon exposure to replication inhibitors [8, 9, 13]. Since our results showed that FA and mTOR pathways work in the same pathway to mediate stalled replication fork restart (Fig. 3), we wanted to determine if mTOR pathway shares the function of FANCD2 in providing fork stability. For this, we have measured IdU tract lengths after 6h exposure to APH. As shown in Figure 4, IdU tract lengths are much longer in the wild type cells as compared to those in the FANCD2-deficient, mTOR-inhibited and FANCD2/mTOR-double deficient cells, strongly suggesting that mTOR is essential for preventing degradation of nascent DNA strands after replication stress. The fact that double-deficient cells did not show any additive effect, further suggests that mTOR and FA pathways are epistatic for this function.

**Fig. 4.**
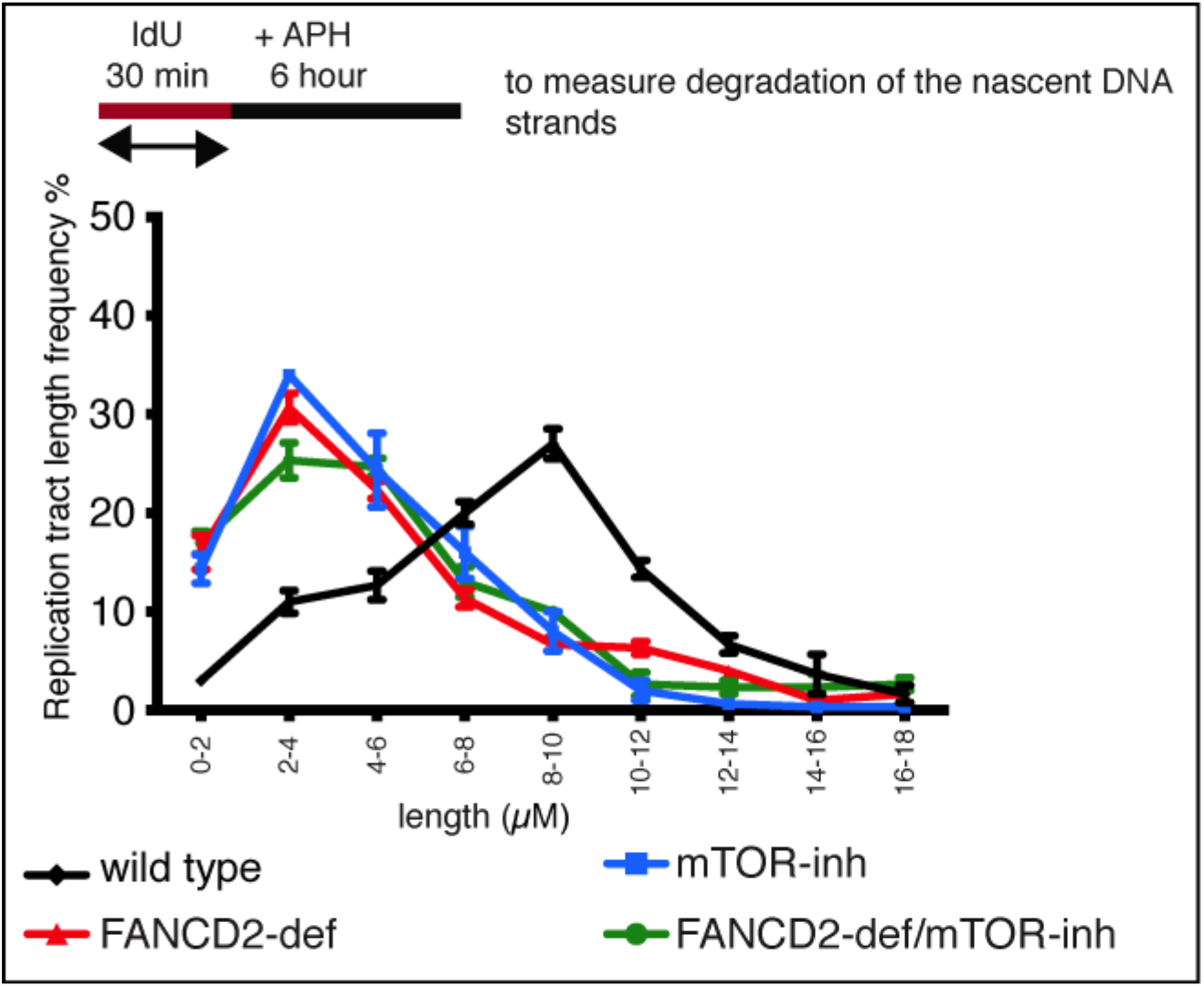
Protection of newly replicated DNA strands after replication fork stalling requires concerted actions of FANCD2 and mTOR. FA patient fibroblast cells PD20 (FANCD2-deficient) and PD20 cells complemented with FANCD2 (PD20+D2, wild type) were treated with mTOR inhibitor rapamycin (20nM) to create FANCD2/mTOR-double-deficient and mTOR-inhibited cells respectively. DNA fiber assay was performed as described in the Fig. 3. To measure degradation of the nascent DNA strands, only IdU (Red colored, first labeling) tract lengths were measured after treatment with APH for 6 hours. Percentage of IdU tract length frequencies were plotted. Compared to the wild type, mTOR-inhibited, FANCD2- single and double-deficient cells showed significantly shorter tracts. Results plotted were from three independent experiments. Error bar represents standard error of mean.

**Fig. 5.**
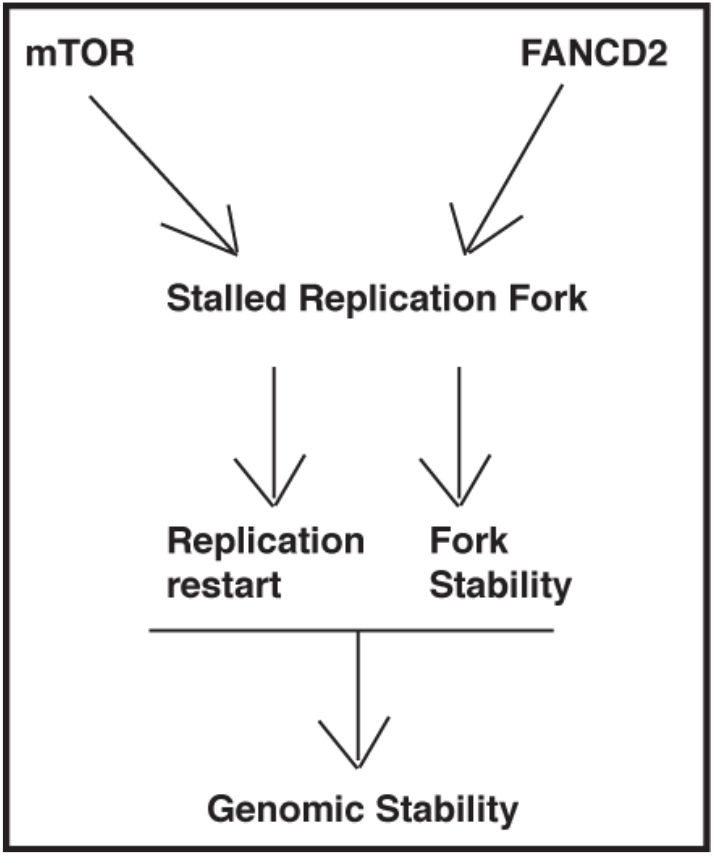
mTOR-FANCD2 model depicting functions at the stalled replication fork. FA protein FANCD2 and mTOR gets associated with each other in response to replication fork stalling. Concerted actions of these proteins mediate stalled replication fork restart, promote fork stability by preventing nascent DNA strands from nucleolytic degradation, and thus contribute to genomic stability.

## DISCUSSION

Several studies indicate that stalled replication fork restart requires concerted actions of a variety of proteins including those involved in the FA DNA repair pathway [8, 9, 11, 12]. FA pathway protein FANCD2 plays a crucial role along with other FA (such as FANCJ, FANCD1) [10] and non-FA (such as Bloom Syndrome protein BLM, FAN1, CtIP) [8, 9, 11, 12] proteins to ensure that replication inhibitor induced stalled forks are again ready to resume and complete the unfinished replication process. Although many proteins contribute to this function, there is a gap in understanding the complete mechanism. Given the involvement of a diverse group of proteins in mediating stalled fork restart, it appears that a much broader and generalized response is involved that regulates multiple steps of the pathway. Although not in the context of replication fork restart, previous studies showed interesting connection between FA and mTOR pathways regulation of DNA damage response [28, 32]. Hence, it is essential to determine if mTOR and FA pathways, specifically via FANCD2, share common functions at the stalled replication forks.

Our results revealed that mTOR and FA pathways share common functions during replication fork stalling to protect cells from replication inhibitor induced stalled forks (Fig. 1A and 1B) but not in response to DNA ICL agent (Fig. 1C). APH and HU are known to induce replication fork stalling [29, 30] whereas MMC causes DNA ICLs [31]. Our result clearly demonstrates a differential response of mTOR pathway to cooperate with the FA pathway, with a functional preference for the stalled replication fork rather than DNA ICLs. This function further becomes evident during DNA damage induced physical association between mTOR and FANCD2. As shown in Fig. 2, while mTOR associates with FANCD2 during fork stalling as well as at the DNA ICLs, this interaction is clearly stronger in response to APH and HU compared to that with MMC (Fig. 2). Although our results clearly suggest a physical interaction between mTOR and FANCD2, it is still possible that a bridging protein mediates this interaction. Future work using recombinant purified proteins will be necessary to determine if this interaction is direct or mediated by another common interactor. DNA ICLs are the hallmark of the FA deficient cells, and functions of FA pathway for repairing ICL has been extensively studied. However, multifunctional FA pathway has an important role during stalled fork repair and restart. Our findings strongly suggest a much broader and widespread association of multiple pathways that are required to mediate efficient fork recovery.

Both APH and HU are replication fork stalling agents [29, 30]. However, they create stalled forks by two distinct means. APH directly inhibits DNA polymerase α [29], whereas HU lowers overall dNTP pool by inhibiting ribonucleotide reductase [30]. Our results show a more prominent role of mTOR within the FA pathway in response to APH rather than HU. mTOR deficiency appears to have a more prominent phenotype in the cellular survival (Fig. 1). More studies will be required to further determine the functional significance of these findings.

Our previous studies showed an essential role of FANCD2 in the stalled replication fork restart. A recent study showed that mTOR is also important for this function. Our results clearly demonstrate that both mTOR and FANCD2 are essential as well as epistatic to ensure efficient fork recovery (Fig. 3). We and others have previously demonstrated another role of FANCD2 at the stalled replication fork. FANCD2 is important to prevent nucleolytic degradation of the nascent DNA strands at the stalled replication fork [8, 9, 13]. Our results demonstrate that mTOR, like FANCD2, protects nascent DNA strands from indiscriminate degradation by the cellular exonucleases in response to the fork stalling agents. Since, FANCD2 and breast cancer associated FA protein BRCA2 (FANCD1) were shown to have concerted role in the stalled replication fork stability [13], future studies are necessary to determine mTOR-BRCA2 interplay at the stalled fork.

Together, our results strongly demonstrate a functional association between two distinct cellular pathways - FA pathway and mTOR pathway during DNA damage induced by the replication inhibitors. Further studies will be necessary to determine the biochemical hierarchy between FANCD2 and mTOR proteins in stalled fork induced chromatin recruitment. Based on this work we propose that stalled fork restart involves more generalized response that stimulates the functional interaction of multiple cell regulatory pathways.

## MATERIALS AND METHODS

### Cell culture

Fanconi anemia patient fibroblast cells PD20 (FANCD2 deficient) and PD20 cells complemented with FANCD2 (PD20+D2) were obtained from FA cell repository at Oregon Health and Science University. Cells were maintained using Dulbecco’s modified Eagle’s medium (DMEM) supplemented with 10% fetal bovine serum (FBS) and penicillin and streptomycin antibiotics at 37°C in 5% CO_2_. MMC, APH and HU were used at different concentrations indicated in the figures. mTOR inhibitor rapamycin was used at 20 nM.

### Cellular viability

Cellular viability was determined by colony formation assay following protocol as described [33]. Wild type (PD20+D2) and FANCD2-deficient (PD20) cells were either untreated or treated with 20 nM rapamycin to create mTOR-inhibited and FANCD2/mTOR double-deficient cells. 500 cells from each cell types were seeded on 60 mm culture dish and were allowed to grow for 8-10 hours. Cells were then treated with different doses of APH, HU or MMC for 6 hours. Next, the cells were grown in fresh media for 7-10 days. Individual colony forming units were counted after staining with 0.1% crystal violet solution.

### Immunoprecipitation and Western blotting

Cells (wild type: PD20+D2 or FANCD2-deficient: PD20) were either untreated or treated with 1 μM MMC for 24h, 1 mM HU for 24h or 30 μM APH for 6 hours. Nuclear extracts were isolated and immunoprecipitation (IP) was performed as previously described [9]. Briefly, nuclear extracts were precleared with IgG and subjected to IP using IgG, anti-mTOR or anti-FANCD2 antibody at 4°C overnight. Samples were then incubated with protein A/G agarose beads for 2h at 4°C. IP was performed in presence of ethidium bromide (10μg/ml), a DNA intercalating agent, to rule out the possibility of DNA-bridged interactions. Beads were then pelleted from the solution, washed three times in IP buffer, boiled in 1X NuPAGE buffer (Life Technologies) and analyzed by SDS-PAGE and Western blotting using specific antibodies against mTOR and FANCD2.

### DNA fiber assay

Single-molecule dual labeling DNA fiber assay was performed as previously described [8, 9, 34] with some modifications. Briefly, cells were treated with 50 μM IdU for 30 minutes to incorporate thymidine analog IdU at the ongoing replicating DNA. Next cells were either untreated or treated with 30 μM APH for 6h to induce replication fork stalling. Then in the fresh media, cells were treated with another thymidine analog CldU for 30 more minutes. Cells were fixed, lysed on silane coated glass slides and allowed to stretch by tilting the slides. Mouse anti-BrdU clone B44 primary antibody (Becton Dickinson, cat # 347580) that targets IdU, and Rat anti-BrdU clone BU1/75 primary antibody (AbD Serotec, cat # OBT0030G) that targets CldU were used to incubate the slides. AlexaFluor conjugated secondary antibodies targeting respective primary antibodies (Thermoscientific, USA) were used to visualize red (IdU) and green (CldU) DNA fibers on a Deltavision microscope (Applied Precision) and analyzed using softWoRx 5.5 software. 50 DNA fibers were counted per sample per experiment and results plotted were from three independent experiments. Error bars represent standard error of means. P values for replication restart experiments were determined from Student’s t-test.

## ACKNOWLEDGMENTS

This work was funded by Faculty Research Enhancement Fund from University of Minnesota, Morris to IC and a grant from the National Institutes of Health R35GM128675 to MKH. We are thankful to Dr. Deanna Koepp at University of Minnesota, Minneapolis for helping with reagents and to Dr. Duncan J. Clarke, University of Minnesota, Minneapolis for providing Deltavision microscope to analyze DNA fiber images. Special thanks to Dr. Rachel Johnson, University of Minnesota, Morris for providing facilities in her research laboratory including biosafety cabinet and CO_2_ incubator and to Prof. Paul Myers, University of Minnesota, Morris for providing facilities in his research laboratory. Authors are also grateful to Prof. Peh Ng., Division Chair, University of Minnesota, Morris for providing all necessary supports that were essential for successful completion of this project. Authors are grateful to Dr. Aileen Bailey and Dr. Rachel Myerowitz, department of Biology, St. Mary’s College of Maryland (IC’s current institution) for providing space and research facilities.

## AUTHOR CONTRIBUTIONS

M.N., M.K.H. AND I.C. designed the study. M.N., K.K. and I.C. performed the experiments and analyzed the data. M.K.H. and I.C wrote the manuscript.

## REFERENCES

1. Niraj, J., A. Farkkila, and A.D. D’Andrea, The Fanconi Anemia Pathway in Cancer. Annu Rev Cancer Biol, 2019. 3: p. 457–478.

2. Adelman, C.A., et al., HELQ promotes RAD51 paralogue-dependent repair to avert germ cell loss and tumorigenesis. Nature, 2013. 502(7471): p. 381–4.

3. Bunting, S.F., et al., BRCA1 functions independently of homologous recombination in DNA interstrand crosslink repair. Mol Cell, 2012. 46(2): p. 125–35.

4. Kim, H. and A.D. D’Andrea, Regulation of DNA cross-link repair by the Fanconi anemia/BRCA pathway. Genes Dev, 2012. 26(13): p. 1393–408.

5. Wang, L.C. and J. Gautier, The Fanconi anemia pathway and ICL repair: implications for cancer therapy. Crit Rev Biochem Mol Biol, 2010. 45(5): p. 424–39.

6. Youds, J.L., L.J. Barber, and S.J. Boulton, C. elegans: a model of Fanconi anemia and ICL repair. Mutat Res, 2009. 668(1-2): p. 103–16.

7. Ceccaldi, R., P. Sarangi, and A.D. D’Andrea, The Fanconi anaemia pathway: new players and new functions. Nat Rev Mol Cell Biol, 2016. 17(6): p. 337–49.

8. Chaudhury, I., et al., FANCD2 regulates BLM complex functions independently of FANCI to promote replication fork recovery. Nucleic Acids Res, 2013. 41(13): p. 6444–59.

9. Chaudhury, I., D.R. Stroik, and A. Sobeck, FANCD2-controlled chromatin access of the Fanconi-associated nuclease FAN1 is crucial for the recovery of stalled replication forks. Mol Cell Biol, 2014. 34(21): p. 3939–54.

10. Raghunandan, M., et al., FANCD2, FANCJ and BRCA2 cooperate to promote replication fork recovery independently of the Fanconi Anemia core complex. Cell Cycle, 2015. 14(3): p. 342–53.

11. Raghunandan, M., et al., Functional crosstalk between the Fanconi anemia and ATRX/DAXX histone chaperone pathways promotes replication fork recovery. Hum Mol Genet, 2019.

12. Yeo, J.E., et al., CtIP mediates replication fork recovery in a FANCD2-regulated manner. Hum Mol Genet, 2014. 23(14): p. 3695–705.

13. Schlacher, K., H. Wu, and M. Jasin, A distinct replication fork protection pathway connects Fanconi anemia tumor suppressors to RAD51-BRCA1/2. Cancer Cell, 2012. 22(1): p. 106–16.

14. Alayev, A. and M.K. Holz, mTOR signaling for biological control and cancer. J Cell Physiol, 2013. 228(8): p. 1658–64.

15. Alayev, A., et al., Effects of combining rapamycin and resveratrol on apoptosis and growth of TSC2-deficient xenograft tumors. Am J Respir Cell Mol Biol, 2015. 53(5): p. 637–46.

16. Alayev, A., et al., Resveratrol prevents rapamycin-induced upregulation of autophagy and selectively induces apoptosis in TSC2-deficient cells. Cell Cycle, 2014. 13(3): p. 371–82.

17. Alayev, A., et al., Estrogen induces RAD51C expression and localization to sites of DNA damage. Cell Cycle, 2016. 15(23): p. 3230–3239.

18. Alayev, A., et al., Combination of Rapamycin and Resveratrol for Treatment of Bladder Cancer. J Cell Physiol, 2017. 232(2): p. 436–446.

19. Holz, M.K., The role of S6K1 in ER-positive breast cancer. Cell Cycle, 2012. 11(17): p. 3159–65.

20. Manna, S. and M.K. Holz, Tamoxifen Action in ER-Negative Breast Cancer. Sign Transduct Insights, 2016. 5: p. 1–7.

21. Yamnik, R.L., et al., S6 kinase 1 regulates estrogen receptor alpha in control of breast cancer cell proliferation. J Biol Chem, 2009. 284(10): p. 6361–9.

22. Alayev, A., S.M. Berger, and M.K. Holz, Resveratrol as a novel treatment for diseases with mTOR pathway hyperactivation. Ann N Y Acad Sci, 2015. 1348(1): p. 116–23.

23. Alayev, A., et al., The combination of rapamycin and resveratrol blocks autophagy and induces apoptosis in breast cancer cells. J Cell Biochem, 2015. 116(3): p. 450–7.

24. Alayev, A., et al., Phosphoproteomics reveals resveratrol-dependent inhibition of Akt/mTORC1/S6K1 signaling. J Proteome Res, 2014. 13(12): p. 5734–42.

25. Berman, A.Y., et al., The therapeutic potential of resveratrol: a review of clinical trials. NPJ Precis Oncol, 2017. 1.

26. Alayev, A., et al., mTORC1 directly phosphorylates and activates ERalpha upon estrogen stimulation. Oncogene, 2016. 35(27): p. 3535–43.

27. Lamm, N., et al., ATR and mTOR regulate F-actin to alter nuclear architecture and repair replication stress, in bioRxiv 2018.

28. Guo, F., et al., mTOR regulates DNA damage response through NF-kappaB-mediated FANCD2 pathway in hematopoietic cells. Leukemia, 2013. 27(10): p. 2040–2046.

29. Ikegami, S., et al., Aphidicolin prevents mitotic cell division by interfering with the activity of DNA polymerase-alpha. Nature, 1978. 275(5679): p. 458–60.

30. Elford, H.L., Effect of hydroxyurea on ribonucleotide reductase. Biochem Biophys Res Commun, 1968. 33(1): p. 129–35.

31. Fujiwara, Y., Defective repair of mitomycin C crosslinks in Fanconi’s anemia and loss in confluent normal human and xeroderma pigmentosum cells. Biochim Biophys Acta, 1982. 699(3): p. 217–25.

32. Guo, F., Mtor-Fanconi Anemia DNA Damage Repair Pathway in Cancer. J Oncobiomarkers, 2014. 2(2).

33. Zhi, G., et al., Fanconi anemia complementation group FANCD2 protein serine 331 phosphorylation is important for fanconi anemia pathway function and BRCA2 interaction. Cancer Res, 2009. 69(22): p. 8775–83.

34. Chaudhury, I. and D.M. Koepp, Degradation of Mrc1 promotes recombination-mediated restart of stalled replication forks. Nucleic Acids Res, 2017. 45(5): p. 2558–2570.

